# Stromal Hedgehog pathway activation by IHH suppresses lung adenocarcinoma growth and metastasis by limiting reactive oxygen species

**DOI:** 10.1101/747915

**Authors:** Sahba Kasiri, Baozhi Chen, Alexandra N. Wilson, Annika Reczek, Simbarashe Mazambani, Jashkaran Gadhvi, Evan Noel, Ummay Marriam, Barbara Mino, Wei Lu, Luc Girard, Luisa M. Solis, Katherine Luby-Phelps, Justin Bishop, Jung-Whan Kim, James Kim

## Abstract

Activation of the Hedgehog (Hh) signaling pathway by mutations within its components drives the growth of several cancers. However, the role of Hh pathway activation in lung cancers has been controversial. We demonstrate that the Hh signaling pathway is activated in lung stroma in a paracrine manner. Genetic deletion of *Shh* in autochthonous murine lung adenocarcinoma had no effect on survival. Early abrogation of the pathway by an anti-SHH/IHH antibody 5E1 led to significantly worse survival with increased tumor and metastatic burden. Loss of IHH by *in vivo* CRISPR led to more aggressive tumor growth suggesting that IHH, not SHH, activates the pathway in stroma to drive its tumor suppressive effects - a novel role for IHH in the lung. Tumors from mice treated with 5E1 had decreased blood vessel density and increased reactive oxygen species (ROS). Treatment of KP mice with 5E1 and N-acetylcysteine, as a ROS scavenger, decreased tumor ROS levels, inhibited tumor growth and prolonged mouse survival suggesting that increased ROS levels from stromal Hh pathway inhibition spurred lung tumor growth. Thus, IHH induces stromal Hh pathway activation to suppress tumor growth and metastases, in part, by limiting ROS production.

## Introduction

Lung cancer is the leading cause of cancer-related mortality in the U.S. and the world (1) with 5-year survival of less than 5% for patients with metastatic disease (2). Non-small cell lung cancer (NSCLC) accounts for ∼85% of lung cancers, of which, adenocarcinoma is the most common subtype of NSCLC (3). *KRAS* mutations are the most common oncogenic driver mutations and occur in ∼30% of lung adenocarcinoma (LAD) (4). Currently, there are no specific targeted therapies available for mutant KRAS LAD.

The Hedgehog (Hh) signaling pathway is critical for embryonic development, tissue homeostasis and cancer (5). The pathway primarily operates in a paracrine manner in which a secreted Hh ligand (SHH, IHH, and DHH in mammals) binds to PTCH, a twelve-pass transmembrane protein, to relieve its basal inhibition of SMO, a seven-pass transmembrane protein. SMO activation leads to activation and nuclear localization of the GLI2 transcription factor to initiate the transcription of target genes, including *PTCH, GLI1,* and *HHIP* (5).

Aberrant activation of the Hh signaling pathway by mutations in pathway components such as *PTCH* (6–11), *SUFU* (11–13), *SMO* (14–18), and amplifications of *GLI1* and *GLI2* (19, 20) drive tumor growth in basal cell carcinoma (BCC) (6–8, 17, 21), medulloblastoma (9, 10, 13, 18–20, 22, 23), keratocystic odontogenic tumors (24, 25), meningioma (12, 14, 16) and ameloblastoma (15). Vismodegib (26) and sonidgeib (27), two potent SMO antagonists, have been approved by the FDA for treatment of locally advanced and metastatic BCC (28, 29). Recently, glasdegib (30), another SMO antagonist, in combination with low dose cytarabine was recently approved by the FDA for acute myeloid leukemia patients 75 years or older or those with comorbidities that preclude intensive chemotherapy (31).

Mutations of Hh pathway components are rare in sporadic epithelial tumors. It was proposed that these cancers recapitulated development by secreting Hh ligands from the tumor epithelia to activate the pathway in stromal cells that, in turn, secreted factors instrumental for tumor initiation and growth (32–35). However, recent data in several epithelial cancers suggest that paracrine activation of stroma by Hh ligands promotes fibroblast expansion and restrains tumor growth early in the tumorigenic process. Inhibition of stromal pathway activation led to accelerated tumor growth with more aggressive, higher grade tumors (36–41).

In lung cancers, a variety of roles for the Hh signaling pathway has been reported. In autochthonous mouse models of small cell lung cancer (SCLC), overexpression of SHH or SMOM2, a constitutively active mutant, in *Rb*^−/−^;*Trp53*^−/−^ cancer cells promoted tumor proliferation (42, 43) and loss of SMO led to significantly decreased tumor formation (42). Chemotherapy-resistant SCLC was more dependent on the pathway for growth and treatment with sonidegib inhibited tumor growth of chemotherapy-resistant SCLC *in vivo* (42). However, a phase III clinical trial showed no benefit of the addition of vismodegib to standard chemotherapy in treatment-naïve SCLC patients (44). For NSCLC, distinct modes of action have been reported for the Hh signaling pathway. In lung squamous cell carcinoma (LSCC) tumor-spheres (45), SOX2 activation induced upregulation of Hh acyltransferase (HHAT) (46), a critical component that palmitoylates Hh ligands (47, 48), and induces autocrine pathway activation to drive growth of LSCC tumor-spheres but not bulk LSCC cells nor lung adenocarcinoma (LAD) tumor-spheres (45). Alternatively, in *PIK3CA*-amplified LSCC, PI3K-mTOR pathway activation led to non-canonical GLI1 expression independent of the Hh pathway (49). GLI1 activation drove LSCC growth and treatment with combinatorial PI3K and GLI1 antagonists leads to tumor regression *in vivo* (49). In LAD tumor-spheres and cell lines, paracrine SHH from LAD epithelia activated the pathway in stroma to express VEGF that in turn, bound to NRP2 receptor to activate the MAPK pathway and express GLI1 in a non-canonical manner (50). Given these varied results of the pathway’s role and modes of action in lung cancers and other solid tumors, we tested the role of paracrine Hh pathway activation in LAD tumorigenesis and growth in autochthonous mutant *Kras^G12D/+^;Trp53^fl/fl^* mouse models of LAD.

## Results

### SHH ligand is expressed in lung adenocarcinoma and activates stromal Hh pathway by a paracrine mechanism

We evaluated the impact of SHH expression on lung adenocarcinoma (LAD) patients as SHH is the primary Hh ligand critical for lung development (51–54), adult lung airway homeostasis (55, 56), and reported in lung cancers (42, 43, 46, 50, 57). We assessed the impact of high *SHH* mRNA expression in LAD patients in the Kaplan-Meier Plotter (KM-Plotter; (58, 59)) database that aggregates Affymetrix microarray mRNA expression with clinical annotation. From 720 LAD patients and using median expression as the cutoff, a univariate Cox regression analysis of high SHH mRNA expression significantly correlated with worse overall survival (*P*= 0.0006; Fig. 1a) and progression free survival (*P*= 0.044; Fig. 1c). These results were corroborated when stage, gender and smoking history were accounted for in multivariate analyses for overall survival (Fig. 1b) but not in progression-free survival (Fig. 1d). We then surveyed 34 human LAD cell lines for Hh ligand expression by qPCR (Fig. 1e). Mutant *KRAS* cell lines were sought as mutant *KRAS* has been reported to upregulate SHH expression (60). The majority of high Hh ligand expressing cell lines, defined as >4x expression of normal bronchial epithelial HBEC7-kt cells, also expressed mutant *KRAS* (Fig. 1e). H23, H2887, HCC44, H2558 LAD cells expressed high levels of SHH protein whereas H441 and H3122 expressed low levels of SHH protein as measured by immunoblot (Fig. 1f), consistent with qPCR results (Fig. 1e).

**Fig. 1.**
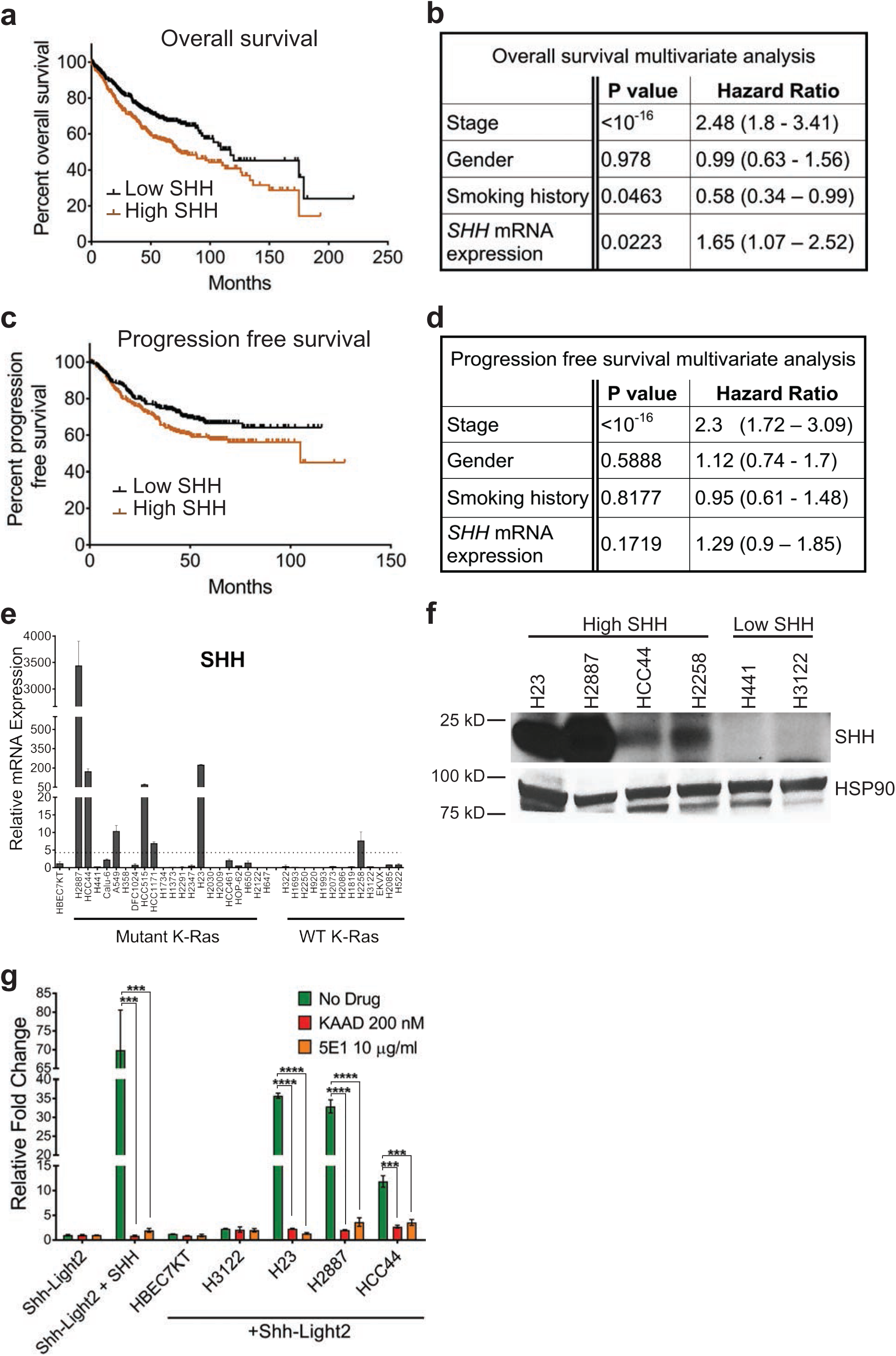
SHH in human lung adenocarcinoma. **a-d** Survival analyses of lung adenocarcinoma patients with high and low *SHH* mRNA expression from Kaplan-Meier Plotter database (58, 59). n=720 patients. High and low mRNA expression is relative to median expression. **a** Kaplan-Meier plot by univariate analysis of overall survival (*P*= 0.0006) is shown. **b** Multivariate analysis of overall survival is shown with stage, gender, smoking history, and *Shh* mRNA expression as variables. **c** Kaplan-Meier plot by univariate analysis (*P*= 0.044) and **d** multivariate analysis of progression free survival of lung adenocarcinoma patients analogous to panels a and b. **e** Expression of *SHH* mRNA as measured by qPCR relative to a normal bronchial epithelial cell line (HBEC7KT). Dashed line represents 4x expression relative to HBEC7KT. **f** Immunoblot of active N-terminal SHH of high and low SHH-expressing lung cancer cell lines from panel e. All qPCR data represent mean of triplicates +/− SD. **g** Relative Hh pathway activity of Shh-Light2 fibroblasts with an 8x-GLI-luciferase reporter is shown. Shh-Light2 cells were co-cultured with low SHH expressing HBEC7KT, a human normal lung epithelial cell line, low SHH expressing H3122 LAD cell line, and high SHH-expressing H23, H2887 and HCC44 LAD cell lines. Cell lines were treated with control vehicle, KAAD-cyclopamine 200 nM, and 5E1 10 μg/ml. *** *P*< 0.001, **** *P*< 0.0001.

To test whether SHH secreted from LAD cells were active, we co-cultured three cell lines with the highest level of SHH (Fig. 1e, f) with Hh-pathway responsive Shh-Light2 mouse embryonic fibroblasts that contain an 8x-GLI binding site-firefly luciferase reporter (61). Treatment of Shh-Light2 cells alone with SHHN conditioned medium (CM) (62) induced high levels of Hh pathway activity that was suppressed by KAAD-cyclopamine 200 nM (63), a potent SMO antagonist, and 5E1 10 μg/ml, a blocking monoclonal antibody that binds to SHH and IHH (64, 65). Co-culture of high SHH-expressing cells (H2887, H23, and HCC44) with Shh-Light2 cells, but without addition of exogenous SHH, resulted in potent activation of the pathway in Shh-Light2 cells, compared to normal airway epithelial HBEC-7kt cells (Fig. 1g). Treatment of these co-cultured cells with KAAD-cyclopamine and 5E1 inhibited Hh pathway activation in Shh-Light2 cells (Fig. 1g). In contrast, low SHH-expressing H3122 cells did not significantly induce Hh pathway activation in Shh-Light2 cells. These results, in particular those with 5E1, suggested that SHH from LAD cells activate the pathway in stromal cells in a paracrine manner.

### SHH does not affect lung adenocarcinoma growth *in vivo*

We next sought to test the role of stromal Hh pathway in lung tumor development. As reliable anti-SHH antibodies for immunohistochemistry (IHC) were not commercially available, we tested for *Shh* mRNA expression by in situ hybridization. We validated *Shh* mRNA probes in the neural tube of E11.5 mouse embryos where SHH is highly expressed in the notochord (66, 67) and floor plate (68, 69) (Supplement Fig. 1). Ten weeks after infection of *Kras^Lox-Stop-Lox-G12D/+^;Trp53^fl/fl^* (*KP*) mice (70) with adenovirus-expressing cre recombinase (adeno-cre) by intranasal inhalation, LAD expressed *Shh* mRNA by in situ hybridization (Fig. 2a). We further verified the expression of *Shh* mRNA specifically *KP* transformed lung epithelia. Lungs from uninfected *KP* mice and *KP;Rosa26^Lox-mtdTomato-Stop-Lox-mGFP/+^* (*KPmTmG*) mice (71), a strain that conditionally switches from constitutive tdTomato expression to GFP expression and initiates LAD when exposed to cre recombinase (Supplement Fig. 2a), infected with adeno-cre were enzymatically dissociated into single cells. Lung epithelial cells were isolated using FACS – CD31- (endothelial cell antigen), CD45- (leukocyte antigen), EpCAM+ (epithelial cell antigen) for uninfected *KP* and EpCAM+, GFP+ (adeno-cre infected cells) for *KPmTmG* epithelia, respectively (Supplement Fig. 2) – and *Shh* mRNA measured by qPCR. Infected *KPmTmG* lung epithelia expressed higher levels of *Shh* mRNA than wild type lung epithelia of uninfected *KP* mice (Fig. 2b). We next tested the requirement of stromal Hh pathway activity for LAD tumorigenesis by crossbreeding *KP* with *Shh^fl/fl^* (72) mice to generate *KP, KP;Shh^fl/+^,*and *KP;Shh^fl/fl^* strains to induce LAD with wild-type (wt), heterozygous, and homozygous loss of SHH expression. Surprisingly, *KP, KP;Shh^fl/+^,*and *KP;Shh^fl/fl^* mice did not show any differences in survival after LAD induction with adeno-cre (Fig. 2c). To verify that *Shh* was indeed deleted in *KP;Shh^fl/fl^* mice, we isolated EpCAM+;GFP+ infected lung epithelial cells by FACS from *KPmTmG* and *KPmTmG;Shh^fl/fl^* mice 3 weeks after adeno-cre infection, analogous to Supplement Fig. 2b, and tested for *Shh* mRNA expression by qPCR. Indeed, *KP;Shh^fl/fl^* infected epithelial cells expressed *Shh* mRNA approximately 6 orders of magnitude less than *KP* epithelial cells (Fig. 2d), suggesting that *Shh* was indeed knocked out. Furthermore, no significant differences in tumor size were seen in the lungs of *KP, KP;Shh^fl/+^,*and *KP;Shh^fl/fl^* 10 weeks after infection (Fig. 2e, f). Taken together, these results suggest that SHH may not play a role in mutant Kras LAD tumorigenesis and progression.

**Fig. 2.**
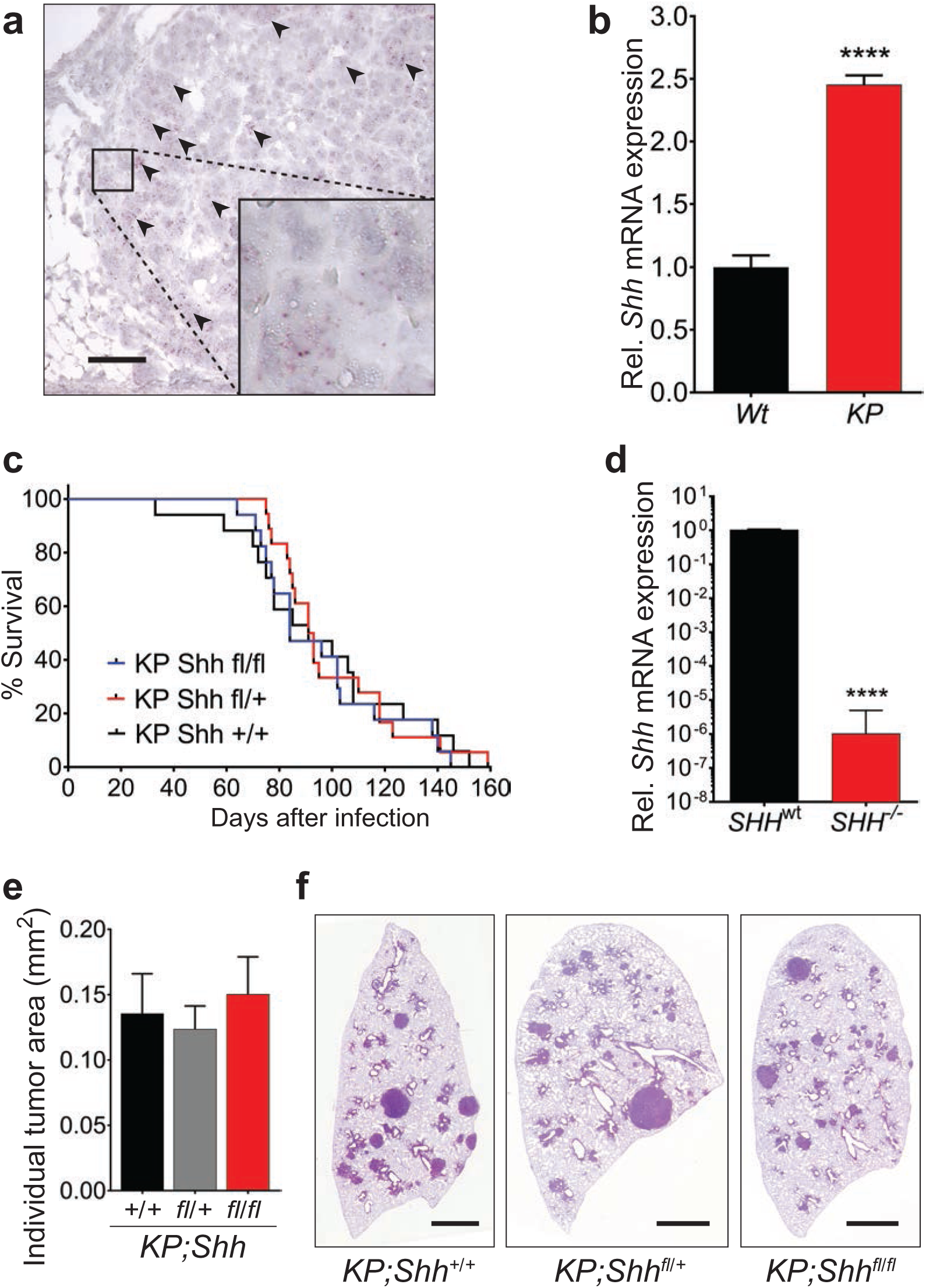
SHH does not affect tumor growth and survival *in vivo*. **a** *Shh* mRNA expression is shown in lung tumor tissues generated in *K-ras^G12D/+^;p53^fl/fl^* (*KP*) mouse by RNA *in situ* hybridization. Red puncta indicate *Shh* mRNA. Scale bar is 50 micrometers. **b** EpCAM+;GFP+ lung epithelial cells of *KPmTmG* mice 3 weeks after adeno-cre recombinase (adeno-cre) infection (‘KP’ in panel) along with CD31-, CD45-, EpCAM+ lung epithelial cells of uninfected *KP* mice (‘Wt’ in panel) were FACS-sorted and *Shh* mRNA levels were analyzed by qPCR. The data represents mean of duplicates +/− s.e.m (n=4 lung lobes from 2 mice) for adeno-cre infected *KPmTmG* mice and mean of triplicates +/− s.e.m (n=4 lung lobes from 4 mice) for uninfected *KP* mice. \*\*\*\**P*< 0.0001. **c** Survival curves of *KP;Shh^+/+^, KP;Shh^fl/+^*, and *KP;Shh^fl/fl^* mice after infection with adeno-cre by intranasal inhalation are shown. *KP;Shh^+/+^* n=17, *KP;Shh^fl/+^* n=18, *KP;Shh^fl/fl^* n=17. **d** EpCAM+;GFP+ lung epithelial cells of *KPmTmG* and *KPmTmG;Shh^fl/fl^* mice 3 weeks after adeno-cre infection were FACS sorted and *Shh* mRNA levels were measured by qPCR. The data represents mean of duplicates +/− s.e.m. n=4 lung lobes from 2 mice per treatment arm. The expression levels were normalized to *Shh^WT^*. \*\*\*\**P*< 0.0001. **e** Quantification of mean tumor area is shown. **f** Representative H&E images of left lung from *KP;Shh^+/+^, KP;Shh^fl/+^*, and *KP;Shh^fl/fl^* mice 10 weeks after adeno-cre infection. Scale bars are 2 millimeters.

### Activation of the Hh pathway in stroma prolongs survival by restraining tumor growth and metastasis *in vivo*

To further examine the effect of paracrine Hh pathway activity in lung tumorigenesis, we first identified 5E1 10 mg/kg twice per week as an optimal dose through an *in vivo* Hh pathway-dependent hair regrowth assay (Supplement Fig. 3) (73, 74). *KP* mice were then treated with 5E1 or IgG_1_ control starting 2 or 6 weeks after tumor initiation by adeno-cre infection (Fig. 3a) such that the pathway was inhibited early in the tumorigenic process (2 weeks) or once adenomas with nuclear atypia had been established (6 weeks) (70). *KP* mice treated with 5E1 starting 2 weeks after tumor initiation had significantly worse survival (Fig. 3b) compared to IgG_1_ treated control mice in contrast to mice treated with 5E1 starting 6 weeks after adeno-cre infection (Fig.3c). Furthermore, *KP* mice treated with 5E1 at the 2 week time point exhibited significantly higher rates of metastases (Fig. 3d), primarily to mediastinal lymph nodes and pleura (Fig. 3 e, f). Examination of LAD tumors after 8 weeks of 5E1 treatment (10 weeks after adeno-cre infection) demonstrated significantly larger tumor burden (Fig. 3g - i) with a greater proportion of poorly-differentiated tumors and less well-differentiated tumors (Fig. 3j, k) compared to mice treated with IgG_1_ control. Thus, pharmacologic inhibition of stromal Hh pathway induced greater tumor burden with greater metastases and worse survival, suggesting that stromal Hh pathway activity restrains LAD growth and metastasis.

**Fig. 3.**
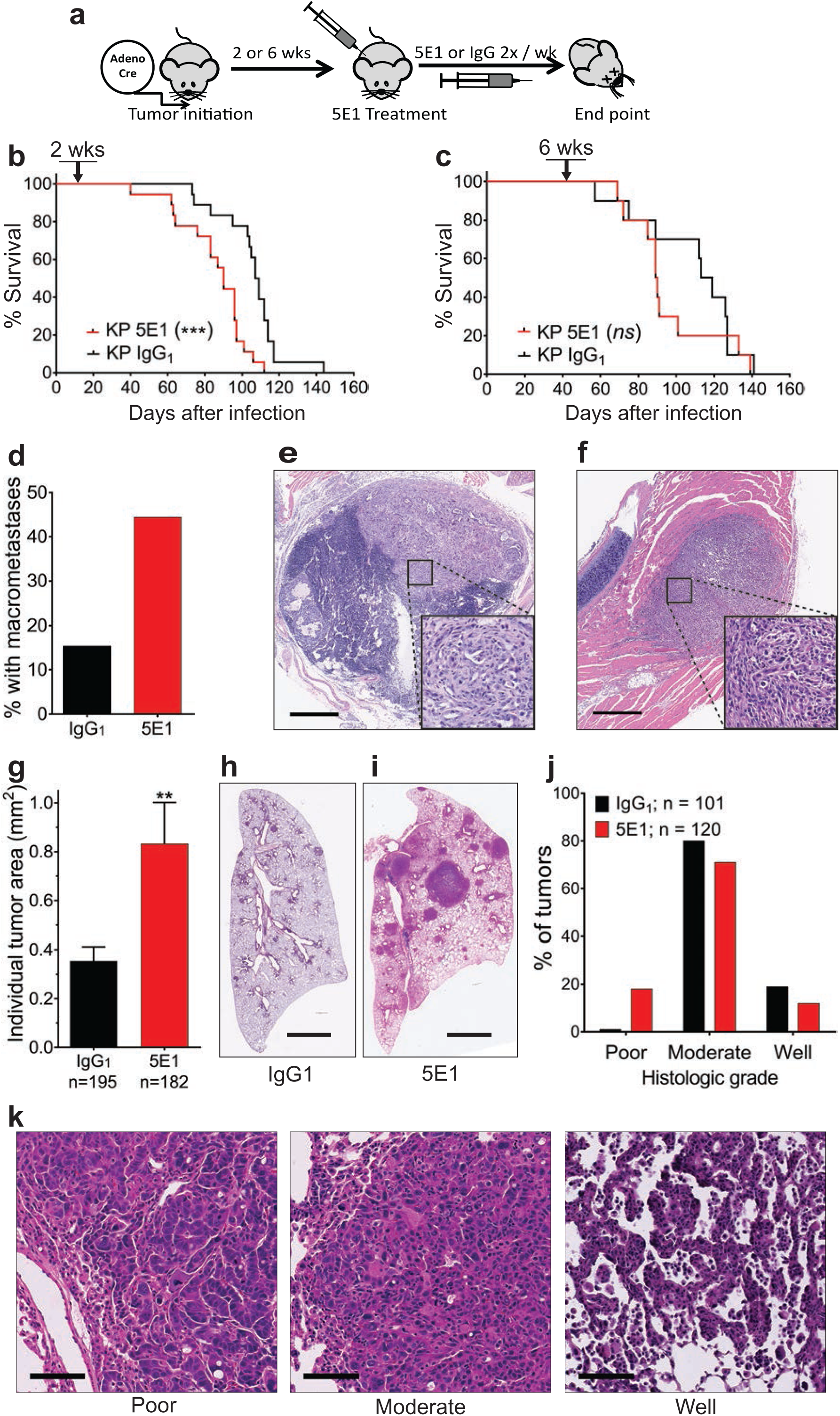
Inhibition of stromal Hh pathway activation worsens survival with increased tumor growth and metastases. **a** Schematic diagram of KP mice infection and treatment with 5E1 or IgG_1._ **b, c** Survival curves are shown of *KP* mice treated with 5E1 10 mg/kg twice per week or corresponding IgG_1_ dose starting **b** 2 weeks (*KP* 5E1 n= 18, *KP* IgG_1_ n=18, \*\*\**P=*0.0002) and **c** 6 weeks (*KP* 5E1 n= 10, *KP* IgG_1_ n=10, *P=*0.34) after tumor initiation by intranasal administration of adeno-cre. **d** Fraction of mice with grossly visible metastases from experiment in panel b is shown. **e, f** Representative H&E images metastatic tumors in a **e** mediastinal lymph node and in **f** pleura invading the chest wall. Scale bars are 500 micrometers. **g-j** *KP* mice were treated with 5E1 10 mg/kg twice per week or corresponding IgG_1_ dose for 8 weeks starting 2 weeks after adeno-cre infection. **g** Quantification of mean tumor area is shown. Data represent mean of IgG_1_ (n = 195 tumors) or 5E1 (n = 182 tumors) +/− s.e.m. \*\**P*< 0.01. **h, i** Representative H&E images of left lung of mice from panel g. Scale bars in panels h and i are 2 and 2.5 millimeters respectively. **j** Percent of tumors with poor, moderate, and well differentiated histologies are shown of *KP* LAD 10 weeks after adeno-cre infection. Data represent mean of IgG_1_ (n = 101 tumors) or 5E1 (n = 120 tumors) +/− s.e.m. **k** Representative H&E images of poor, moderate, and well differentiated tumors are shown. Scale bars are 100 micrometers.

### IHH is the predominant Hh ligand in murine mutant Kras LAD

With the disparate outcomes of genetic SHH loss (Fig. 2c, e, f) and pharmacologic blockade by 5E1 (Fig. 3b, d - k), we hypothesized that IHH may play a role in LAD tumorigenesis as 5E1 binds both SHH and IHH. We verified that 5E1 can inhibit stromal pathway activation by IHH using Shh-Light2 cells stimulated with either recombinant IHH (rIHH) or rSHH (Fig. 4a). Of note, there was almost no induction of pathway activity with recombinant DHH treatment (results are not shown). As reliable antibodies for IHH IHC and immunoblots were not commercially available, we turned to RNA *in situ* hybridization. *KP* LAD 10 weeks after adeno-cre infection expressed substantial amounts of *Ihh* mRNA (Fig. 4b). To further verify that IHH is expressed by transformed lung epithelial cells, EpCAM+,GFP+ epithelial cells were isolated by FACS (analogous to Supplement Fig. 2b) from *KPmTmG* mice 6 weeks after adeno-cre infection. The sorted epithelial cells show a striking increase in *Ihh* mRNA expression in comparison to *Shh* mRNA as measured by qPCR (Fig. 4c). To genetically test the requirement of IHH to suppress LAD tumorigenesis and growth, we used the pSECC lentiviral *in vivo* CRISPR/Cas9 system (75) that encodes for Cre recombinase to initiate tumorigenesis, Cas9 for gene editing, and sgRNA against the gene of interest. Several candidate sgRNA against *Ihh* (sg*Ihh*) were tested with SURVEYOR assay (Supplement Fig. 4) and the sgRNA sequence (#2, hereafter just sg*Ihh*) with the greatest percentage of digested bands was chosen for further study. We tested pSECC-*Ihh* for loss of *Ihh* mRNA expression by qPCR in 808-T3 cells, a murine *KP* LAD cell line with high *Ihh* mRNA expression (Supplement Fig. 5a). Approximately half of the clones from 808-T3 cell lines transfected with the pSECC-*Ihh* showed substantial decreases in *Ihh* mRNA expression compared to pSECC-*GFP* control (Supplement Fig. 5b). Subsequently, *KP;Rosa26^LSL-fLuc/+^* (*KPL)* mice were infected with lentiviral particles containing pSECC-*Ihh* or pSECC-*GFP* via intratracheal administration and tumor growth monitored by bioluminescence imaging (BLI). Infection with pSECC-*Ihh* induced significant tumor growth compared to pSECC-*GFP* control 18 weeks after infection (Fig. 4, d-f). *KPL* mice infected with pSECC-*GFP* eventually developed tumors that were detected by BLI at 22-26 weeks after infection (Fig. 4d).

**Fig. 4.**
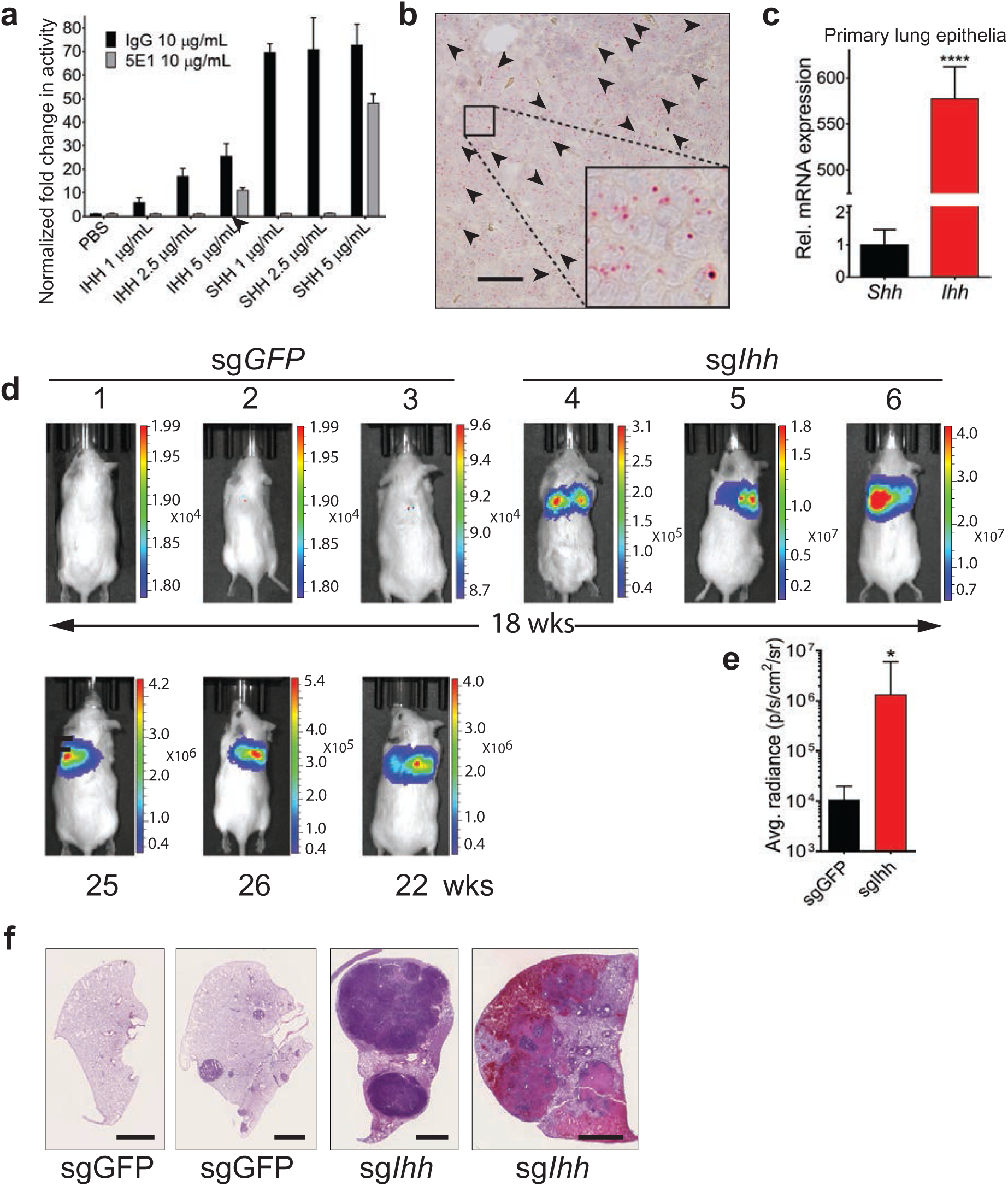
*IHH* regulates the suppression of lung adenocarcinoma. **a** Hh pathway activity as measured by 8x-GLI-luciferase reporter relative to PBS control in Shh-Light2 reporter cells is shown. Shh-Light2 cells were treated with 1, 2.5, and 5 µg/mL of mouse rIHH or rSHH in combination with 5E1 or IgG_1_ 10 µg/mL. PBS was used as a control vehicle for IHH or SHH. **b** In situ hybridization for *Ihh* mRNA in LAD of *KP* mice is shown. Red puncta indicate *Ihh* mRNA. Scale bar is 50 micrometers. **c** *Shh* and *Ihh* mRNA levels of FACS sorted lung epithelial cells from *KPmTmG* mice 6 weeks after adeno-cre infection is shown. mRNA was measured with qPCR. Data are mean of duplicate +/− s.e.m. n=4 mice. \*\*\*\**P*< 0.0001 **d** Bioluminescence images (BLI) are shown of lung tumors in *KP;Rosa26^LSL-fLuc/+^* 18 weeks after infection with lentiviral pSECC-sg*Ihh or* pSECC-sgGFP. The second row shows BLI of mice 22-26 weeks after pSECC-sgGFP infection. **e** Quantification of luminescence intensity is shown for pSECC-sg*Ihh* and pSECC-sgGFP infected mice at 18 weeks. **f** Representative images of H&E images of right upper lobes of pSECC-sg*Ihh* and pSECC-sgGFP infected mice at 18 weeks. Scale bars are 2 millimeters.

### IHH in human lung adenocarcinoma

We next tested for *IHH* mRNA by in situ hybridization in human LAD samples in mutant and wild type *KRAS* and *TP53* samples. Two of the three mutant *KRAS;TP53* samples expressed *IHH* mRNA in malignant cells whereas only one of the six wild type samples expressed *IHH* mRNA (Fig. 5a, Supplement Fig. 6). All of the *IHH* mRNA positive tumors had a predominance of lepidic histology with mucinous features (Supplement Fig. 6). Lepidic histology has been correlated with less aggressive biology. The prognosis of mucinous histology in LAD is uncertain currently (76). Re-examination of the 34 human LAD cell lines (Fig. 1e) revealed only 4 lines with *IHH* mRNA significantly elevated beyond the normal lung epithelial line HBEC-7kt (Fig. 5b). As most of the cell lines were generated from patients with late stage or metastatic adenocarcinomas, the dearth of cancer lines with upregulated *IHH* mRNA corroborates the in situ results of *IHH* mRNA in more indolent lepidic histologies (Fig. 5a, Supplement Fig. 6). Univariate Cox regression analysis of a clinically annotated microarray database of human LAD (KM Plotter; (58)) revealed that patients with high expression of *Ihh* mRNA had better overall (*P* = 0.0001; Fig. 5c) and progression free (*P* = 0.0069; Fig 5e) survival compared to those with low expression. These results remain consistent after multivariate analyses when stage, gender, and smoking history are considered (Fig. 5d, f), in agreement with our murine LAD data (Fig. 3b and 4, d-f). The data here suggest that IHH is sufficient to suppress tumor initiation and growth and that SHH is dispensable for LAD tumorigenesis.

**Fig. 5.**
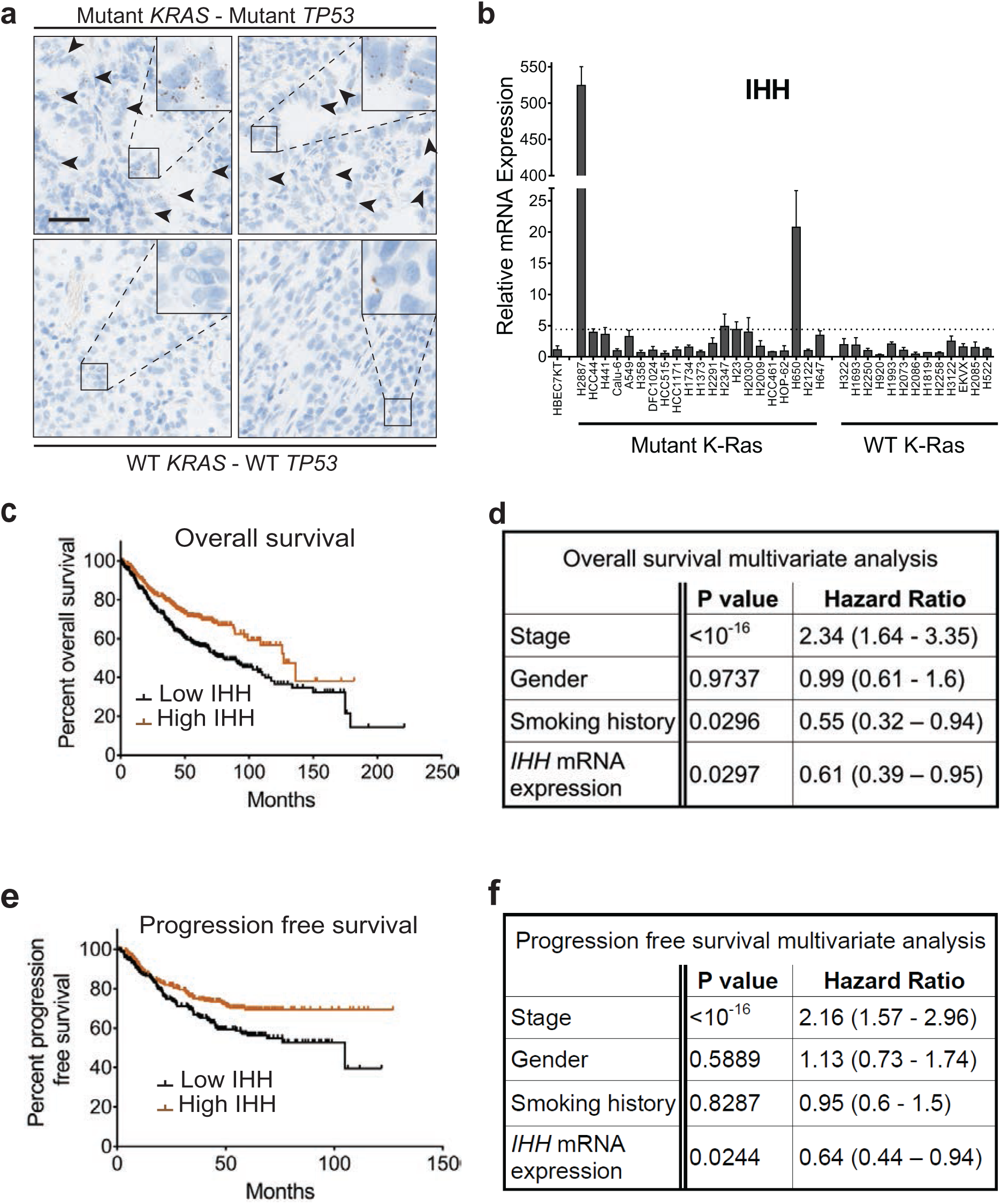
IHH in human lung adenocarcinoma. **a** In situ hybridization for *IHH* mRNA in human LAD is shown. Brown puncta indicate *IHH* mRNA. Arrowheads indicate areas of *IHH* mRNA staining in malignant cells. Scale bar is 50 micrometers. **b** Expression of *IHH* mRNA as measured by qPCR relative to a normal bronchial epithelial cell line (HBEC7KT). Dashed line represents 4x expression relative to HBEC7KT. **c-e** Survival analyses of lung adenocarcinoma patients with high and low *IHH* mRNA expression from Kaplan-Meier Plotter database (58, 59). n=673 patients. High and low mRNA expression is relative to median expression. **c** Kaplan-Meier plot by univariate analysis of overall survival (*P*= 0. 0001) is shown. **d** Multivariate analysis of overall survival is shown with stage, gender, smoking history, and *Ihh* mRNA expression as variables. **e** Kaplan-Meier plot by univariate analysis (*P*= 0. 0069) and **f** multivariate analysis of progression free survival of lung adenocarcinoma patients analogous to panels c and d.

### Loss of Stromal Hh pathway inhibits angiogenesis and increases reactive oxygen species

The Hh signaling pathway has been implicated in the regulation of angiogenesis in normal tissues (77, 78) and cancer (79, 80) through induction of angiogenic factors including VEGFs and ANG1, 2. Examination of CD31 expression, a marker of endothelial cells, showed decreased blood vessel density in LAD tumors treated with anti-SHH/IHH 5E1 antibody compared to IgG_1_ treated tumors (Fig. 6a, b). As the effects of stromal Hh pathway inhibition were seen with mice when treatment was initiated 2 weeks after adeno-cre infection (Fig. 3b, d-k), we hypothesized that the inability of growing tumors to generate new vessels would lead to early hypoxia and production of reactive oxygen species (81, 82), that in turn, would promote tumor proliferation and growth (83–85). We developed 2 macros (*ROI_Draw* and *Nuclear_Fraction_Calculator*) for ImageJ (86) or Fiji (87) to quantify DAB stained nuclei of phospho-histone 2AX (γH2AX), a protein that responds to double stranded DNA breaks and a marker of oxidative stress (88, 89). *Nuclear_Fraction_Calculator* counts DAB stained nuclei and total nuclei in digital images of tissue sections and calculates the fraction of IHC positive nuclei within regions of interest (ROI; tumors in our studies) that have been drawn interactively with *ROI_Draw.* With these macros, LAD from mice treated with 5E1 showed significantly higher fraction of nuclei stained with γH2AX than tumors from IgG_1_ treated mice (Fig 6c, d), suggesting increased DNA damage from reactive oxygen species (ROS). To assess whether ROS from stromal Hh pathway inhibition induced accelerated tumor growth, *KP* mice were treated with 5E1 and *N-*acetyl cysteine (NAC), as a scavenger of ROS and precursor to the antioxidant, glutathione (GSH) (Fig. 6e). Treatment with NAC and 5E1 prolonged survival compared to 5E1 and vehicle control (Fig. 6f) whereas treatment with NAC and IgG_1_ did not affect survival (Fig. 6g). Furthermore, the median survival of 5E1 with NAC approximated that of IgG_1_ with vehicle control (Supplement Fig. 7). Interestingly, the rate of metastases did not decrease when mice were treated with 5E1 and NAC compared to 5E1 and vehicle control (Fig. 6h). However, the tumor burden of mice treated with 5E1 and NAC decreased substantially compared to mice treated with 5E1 and vehicle control 10 weeks after adeno-cre infection (Fig. 6i, j) and corresponded to decreased ROS as measured by γH2AX stained nuclei (Fig. 6k, l). These data suggest that IHH restrains tumor growth through support of angiogenesis with limiting ROS production early in the tumorigenic process.

**Fig. 6.**
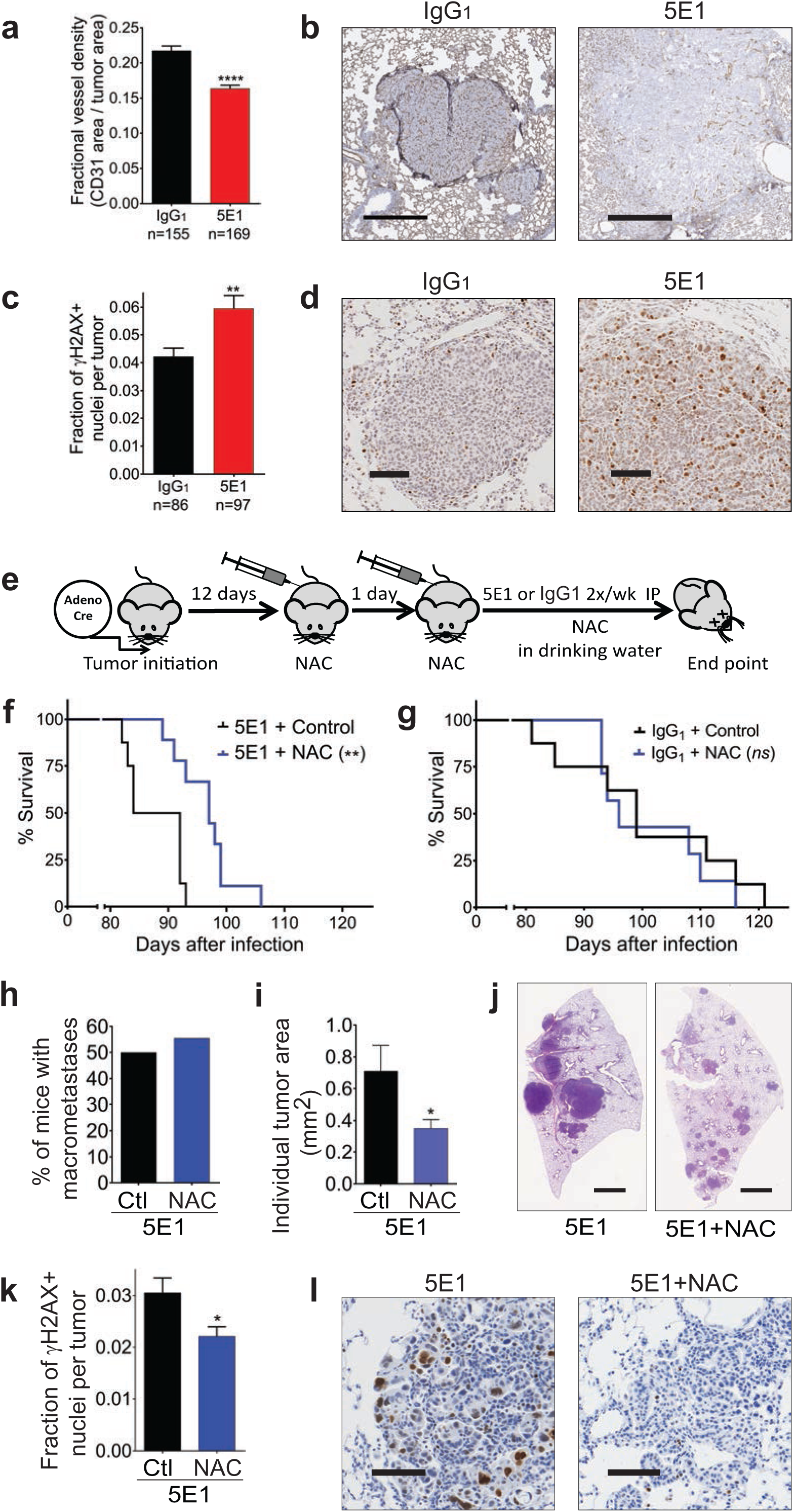
*IHH* loss inhibits angiogenesis and increases reactive oxygen species. **a-d** *KP* mice were treated with 5E1 10 mg/kg twice per week or corresponding IgG_1_ dose for 8 weeks starting 2 weeks after infection **a** Quantification of vessel density (area of CD31 positive cells in tumor /tumor area) is shown. Data represent mean of IgG_1_ (n=155 tumors) or 5E1 (n=169 tumors) +/− s.e.m. \*\*\*\**P*< 0.0001. **b** Representative images of lung tumors of *KP* mice stained for CD31 by IHC with DAB substrate. Scale bar is 500 micrometers. **c** Fraction of γH2AX+ nuclei (γH2AX+ nuceli per tumor/total nuclei per tumor) is shown. Data represent the mean of IgG_1_ (n = 86 tumors) or 5E1 (n = 97 tumors) +/− s.e.m. \*\**P*< 0.01 **d** Representative images of lung tumors of *KP* mice stained for γH2AX by IHC. DAB was used as substrate. Scale bar is 100 micrometers **e** Schematic diagram of survival study for panels f-h. *KP* mice were infected with adeno-cre by intranasal inhalation and treated with vehicle or N-Acetyl Cysteine (NAC) 200mg/kg i.p. once per day on days 12 and 13 after adeno-cre infection. From day 14, mice were treated with 5E1 10 mg/kg i.p. twice per week or corresponding IgG_1_ dose. and NAC 1g/L supplemented in their drinking water. **f** Survival curves are shown of *KP* mice treated with 5E1 and control vehicle (n= 8) or NAC (n=9). \*\**P=*0.0031. **g** Survival curves of *KP* mice treated with IgG_1_ with control vehicle (n= 8) or NAC (n=7, *P =* 0.55). starting 2 weeks after infection. **h** Fraction of mice with grossly visible metastases from experiment in panel f is shown. **i** Quantification of mean lung tumor area of *KP* mice treated with 5E1 in combination with control or NAC for 8 weeks starting 2 weeks after adeno-cre infection.. Data represent the mean of Ctl (n = 64 tumors) and NAC (n = 85 tumors) +/- s.e.m. \**P*< 0.05. **j** Representative H&E images of left lung from panel i. Scale bars are 2 millimeters. **k** Fraction of γH2AX+ nuclei of tumors is shown. Data represent the mean of Ctl (n = 87 tumors) and NAC (n = 82 tumors) +/− s.e.m. \**P*< 0.05. **l** Representative images of lung tumors of *KP* mice from panel k stained for γH2AX by IHC. DAB was used as substrate. Scale bars are 100 micrometers.

## Discussion

We have shown that mutant KRAS LAD secretes Hh ligands to activate stromal cells in a paracrine manner (Fig. 1g), that IHH (Fig. 3), rather than SHH, is the dominant ligand in early transformed lung airway cells and that it restrains tumor growth (Fig. 3g-i, 4d-f), grade (Fig. 3j, k) and metastases (Fig. 3d-f). Inhibition of stromal Hh pathway activation by IHH decreased angiogenesis (Fig. 6a, b) and significantly increased ROS (Fig 6c, d) leading to rapid tumor growth (Fig. 6i, j) and shorter survival (Fig. 6f).

In accordance with previous studies (36–40, 90), paracrine Hh activation of stroma, particularly early in the tumorigenic process, suppresses lung tumor growth, formation of aggressive histologies and metastases. A surprising result of our studies was the central role of IHH, instead of SHH, to suppress tumor growth. SHH is the dominant ligand that regulates lung development (51–54, 91), adult lung airway homeostasis (55, 56, 92), and lung cancers (42, 43, 46, 50, 57). IHH is expressed in the adult colon (93) and prostate (90) and restrains the growth of colon (40, 94) and prostate (41) cancers. However, to our knowledge, IHH activity has not been reported in the lung. Further studies are needed to test if IHH has a role in the homeostasis of the adult lung epithelia or if it is unique to lung cancers.

In our studies, loss of stromal pathway activation in *KP* LAD decreased blood vessel density (Fig. 6a, b) suggesting that the Hh signaling pathway induces angiogenesis in the lungs consistent with reports in other organs (77, 78, 95). However, loss of stromal Hh pathway activation in *KP* pancreas ductal adenocarcinoma (PDAC) had the opposite effect and led to increased tumor blood vessel density (35, 36). Inhibition of angiogenesis through VEGFR2 antagonism in *KP;Shh^fl/fl^* PDACs prolonged mouse survival (36). Another study reported that loss of Hh ligand co-receptors, GAS1 and BOC, in mouse embryonic and pancreas cancer-associated fibroblasts (CAFs) led to partial suppression of pathway response to SHH and increased angiogenesis (96). Loss of co-receptors GAS1, BOC, and CDO in fibroblasts caused a more severe suppression of the pathway and inhibited angiogenesis through modulation of angiogenic ligands VEGFA, ANGPT1, 2 (96). If stromal cells respond distinctly to SHH and IHH ligands, then IHH may play a more prominent role in angiogenesis in LAD than SHH due to the lower potency of IHH (Fig. 4a) analogous to the diminished pathway response of *Gas1*^−/−^;*Boc*^−/−^ fibroblasts in pancreatic cancer (96). Previous studies also have noted the differences in genomic and transcriptomic heterogeneity (97) and effectors downstream of mutant Kras (98) between murine *KP* LADs and PDACs. Such differences may also play a role in the tumor microenvironment where responses to Hh ligands may differ significantly between pancreas and lung stroma. The distinct phenotypic outcomes of stromal Hh pathway activation in LAD and PDAC suggest that tumor-stromal interactions of various cancer types will need to be studied individually and cautions against broad generalizations.

ROS exhibit seemingly paradoxical effects of tumor growth enhancement and tumor cytotoxicity depending on their levels (99). Oncoproteins, such as mutant KRAS and MYC, and hypoxic states can increase cellular ROS levels (85, 100) that enhances tumor growth (85, 101–103). But high levels of ROS can be cytotoxic and cancer cells upregulate antioxidant proteins including glutathione peroxidases, peroxiredoxins, and NRF2 to maintain ROS at optimal levels (100). Here, we have shown that loss of stromal pathway activity early in the tumorigenic process increased ROS in tumor cells (Fig. 6c, d). Reduction of ROS with NAC combined with stromal pathway inhibition prolonged survival with retardation of tumor growth in *KP* LAD (Fig. 6f, i-l). Our results suggest that *KP* LAD can tolerate higher levels of ROS to accelerate tumor growth. Further studies will be needed to identify the upper limit of tolerable ROS levels before cytotoxic effects become dominant.

Our studies here highlight the tumor suppressive roles of stromal Hh pathway activation by IHH via limiting hypoxia and ROS generation through angiogenesis and reinforce the anti-oncogenic role of stroma early in the tumorigenic process. Identification of factors that negatively regulate IHH production in LAD may serve as targets of small molecule or antibody therapeutics to enhance IHH expression and restrain tumor growth and metastases. Such therapeutic strategies may be employed in in early stage or locally advanced disease prior to surgery/high dose radiation or concurrent chemoradiation, respectively, where treatment failure often occurs due to distant metastases. Also, identification of such factors may serve as biomarkers to determine the early stage patients that might benefit from more aggressive therapy.

## Materials And Methods

### Cell culture

All human lung adenocarcinoma cell lines were obtained from the Hamon Cancer Center Collection (UT Southwestern Medical Center, UTSW), were DNA fingerprinted with a PowerPlex 1.2 kit (Promega) and tested for mycoplasma using e-Myco kit (Boca Scientific). The cell lines were generated between 1979 and 2007. Cells were maintained in RPMI-1640 (Life Technologies) with 5% fetal bovine serum (FBS). 808-T3 and Green-Go (75) cell lines were kind gifts from Dr. David McFadden (UTSW) and Dr. Tyler Jacks (MIT), respectively, and were maintained in DMEM (Life Technologies) with 10% FBS. All cells were maintained at 37°C, with 5% CO2, and under humidified conditions.

### Drugs and reagents

5E1 antibody was generated in our laboratory (see supplemental material and methods) and prepared in PBS. IgG_1_ (InVivoMab, BE0083) was diluted in PBS. KAAD-cyclopamine (Millipore) was prepared in DMSO. Recombinant SHH (C25II) (R&D Systems) and IHH (C28II) (Genscript) were prepared in PBS containing 0.1% bovine serum albumin (BSA). N-Acetyl-L-cysteine (NAC) was purchased from Sigma-Aldrich and prepared in PBS for i.p. injection or sterile tap water for supplemented drinking water. For NAC solution, pH was adjusted to 7.4.

### GLI-reporter assay

Shh-Light2 cells (61), a clonal NIH-3T3 cell line that stably expresses 8xGLI-binding site-firefly and TK-*Renilla* luciferase reporters, were co-cultured with LAD cell lines in 24-well plates until confluent and then treated with KAAD-cyclopamine (Millipore) 200nM, 5E1 antibody 10 µg/ml or recombinant SHHN protein 1µg/ml in DMEM containing 0.5% (vol/vol) bovine calf serum. Luciferase activity was measured by Fluostar Optima (BMG Labtech) using Dual Luciferase Assay Reporter System (Promega).

### Quantitative real-time PCR

Total RNA was extracted using TriZol (Invitrogen) and then purified with PureLink RNA Mini Kit (Invitrogen). cDNA was generated using iScript cDNA synthesis kit (Bio-Rad) or Superscript III First Strand Synthesis System (Invitrogen). qPCR was performed using Bio-Rad CFX real-time cycler and SYBR Green Master Mix (Bio-Rad). Data are presented as fold change relative to control samples using the ΔΔCt (2^−ΔΔCt^) method with *HPRT1 or GAPDH* as an internal control gene. Primers for qPCR are listed in Supplementary Table 1.

### Western blot

Cell lysates were generated and analyzed as previously described (104). Briefly, cells were lysed in ice-cold lysis buffer (M-PER Mammalian Protein Extraction Reagent (Thermo Scientific) with protease inhibitors (Roche) and PhosSTOP phosphatase inhibitors (Roche). Cell lysates were centrifuged at 14,000 rpm for 5 min at 4°C and then supernatants were collected. Protein concentration was measured using BCA protein assay kit (Pierce) following the manufacturer’s instructions. The following primary antibodies were used: SHHN (1:1000, Cell Signaling Technology, C9C5) and HSP90 (1:2000, Santa Cruz biotechnology, sc-13119).

### sg-RNA design and cloning

All sg-RNA against *Ihh* were designed using GE Dharmacon web tool. The sg-RNA sequences targeting GFP was published previously (105). sg-RNA oligo candidates (listed on supplementary table 2) were inserted into pSECC vector (a kind gift from Dr. Tyler Jacks, Addgene, 60820) by following the protocol available at this website: http://genome-engineering.org/gecko/.

### Co-transfection of 808-T3 cells

Cells were grown to 70% confluency on 6 well plates and then co transfected with pCMV:DsRed(FRT)GFP plasmid expressing DsRed (Addgene, 31128) and pSECC-*Ihh* or pSECC-*GFP* using Lipofectamine 3000 (Thermo Fisher Scientific) following manufacturer instructions. DsRed+ transfected cells were FACS sorted and plated at limiting dilutions to isolate clonal lines.

### Animals

All animal related experiments and procedures were performed with prior approval of the Institutional Animal Care and Use Committee at UTSW. FVB, *Kras^Lox-Stop-Lox-G12D/+^, Trp53^fl/fl^, Shh^fl/fl^* and, *Rosa26^Lox-mtdTomato-Stop-Lox-mGFP/+^* mice were purchased from Jackson Laboratory (Bar Harbor, ME) and compound strains were generated through cross-breeding.

### Infection and Treatment of Mice

Adenovirus-expressing cre recombinase (Ad5-CMV-Cre) was purchased from Vector Development Laboratory (Baylor College of Medicine, Houston). Six to ten weeks old mice were infected by intranasal instillation with 3 x 10^8^ pfu per mouse as described previously (106) to initiate lung tumorigenesis. For the *in vivo* CRISPR experiments, 10 – 14 week old *Kras^Lox-Stop-Lox-G12D/+^; Trp53^fl/fl^;Rosa26^LSL-fLuc/+^* (*KPLuc*) mice were infected with 5×10^4^ ifu of lentivirus containing pSECC-*Ihh* or pSECC-GFP via intratracheal administration as described previously (106). *KP* or *KP;Rosa26^Lox-mtdTomato-Stop-Lox-mGFP/+^ (KPmTmG)* mice were treated with 5E1 or IgG_1_, 10 mg/kg intraperitoneally (i.p.) twice per week starting 2 or 6 weeks after adeno-cre infection. For NAC study, *KP* mice were infected with adeno-cre then treated with NAC 200 mg/kg i.p. on days 12 and 13 after adeno-cre infection. Afterwards, NAC 1 g/L (pH=7.4) was provided in the drinking water. Supplemented drinking water was changed every 2-3 days for the duration of study.

### Lung tissue extraction and processing

Mice were anesthetized with Avertin 25 mg/kg i.p., lungs perfused with ice-cold PBS, inflated with ice-cold 4% Paraformaldehyde (PFA) in PBS by intra-tracheal instillation, then fixed in 4% PFA at 4°C for 24 hours. Tissue processing and paraffin embedding were performed by Tissue Management Core Facility or Histo-Pathology Core Facility at UTSW. Frozen lung tissue blocks were made by inflating lungs with 50% (v/v) OCT (Tissue-Tek) in PBS and embedded in cryomold with 100% OCT on dry ice, and stored in −80°C. Five and fifteen micron thick sections were made from each PFA fixed paraffin-embedded and frozen tissue blocks respectively and subjected to hematoxylin and eosin (H&E) or immunohistochemistry staining. Brightfield images were taken using a Nikon Eclipse E800 or Hamamatsu Nanozoomer in Whole Brain Microscopy Facility (UTSW). Tumor area on H&E stained images were measured using NIS Elements imaging software (Nikon). The fraction of IHC positive nuclei in each tumor was estimated using ImageJ or Fiji as described in supplemental material and methods.

### Immunohistochemistry

Heat-mediated antigen retrieval (citrate buffer, pH 6) was used for tissue sections from paraffin-embedded blocks. Goat serum (Sigma) or donkey serum (Sigma) was used to block for 1 hour, and diluted primary antibodies were applied at 4°C overnight. Vectastain ABC (Vector Labs) with DAB substrate (Vector Labs) was used to optimize staining according to the manufacture’s protocol. The following primary antibodies were used: Ser139-p-Histone H2A.X (1:1,000; Cell Signaling Technology, 9718) and CD31(1:500, Cell Signaling Technology, 77699).

### RNA in situ hybridization method (RNAScope)

#### Murine Samples

Five micrometer sections from paraffin embedded lungs were deparaffinaized, fixed in 10% formalin solution at room temperature for 24 hours and then subjected to RNAscope assay using RNAscope 2.0 HD Reagent Kit-Red (Advanced Cell Diagnostics (ACD), 310034) and following manufacturer instructions. Mm-Ihh-noXHs (413091) and Mm-Shh (314361) probes were used for murine *Ihh* and *Shh* mRNA detection, respectively. Dapb (negative control, 310043, ACD) and PPIB (positive control, 313911, ACD) were used for quality control (data not shown).

#### Human Samples

Please see Supplementary Methods for full details. Briefly, in situ hydbridization was performed on an automated Leica Bond RX autostainer (Leica Biosystems, Nussloch, GmbH). LS 2.5 Probe-Hs-IHH probe (472388, ACD) was used. RNA expression of IHH was scored using a semi-quantitative scoring system as follows: 0: no staining or <1 dot/10 cells; 1+: 1-3 dots/cell; 2+: 4-9 dots per cell, None or very few dot clusters; 3+: 10-15 dots/ cell and <10% dots are in clusters; 4+: >15 dots/cell and >10% dots are in clusters. Positive (PPIB, Hs-PPIB, 313908) and negative (Dapb, Hs-PPIB, 312038) control probes were also evaluated, dapB score of <1 and PPIB score ≥2 with relatively uniform PPIB signals throughout the sample were considered adequate for analysis (data not shown).

### Histology Analysis

H&E stained lungs with tumors from *KP* mice were examined. The pathologist was blinded to the conditions of the experiment. As nearly all tumors <0.5 mm were well-differentiated histology, only tumors ≥ 0.5 mm were examined. Tumors were graded as poor, moderate or well differentiated cancers.

### Digestion of lung tissue and FACS-sort of lung epithelial cells

Single cell suspensions of whole lungs were prepared as described previously (107). For FACS, single cell suspensions were incubated with eBiosccience Fixable Viability Dye eFluoor™ 780 (Invitrogen) and the following antibodies (0.6 μg per 10**^7^** cells): PerCP-Cy5.5 Rat Anti-Mouse CD45 (BD Pharmingen, 550994), PE-Cy7 Rat Anti-Mouse CD31 (BD Pharmingen, 561410), and Brilliant Violet 421 anti-mouse CD326 (Ep-CAM) (BioLegend, 118225) on ice for 45 minutes, and then subjected to FACS-sorting using FACS Aria II (BD Biosciences) at the Moody Foundation Flow Cytometry Core Facility at the Children’s Research Institute at UTSW. Flow cytometry data were analyzed with FlowJo v10.

### Statistical analysis

GraphPad Prism 7 software was used to generate the graphs and for statistical analysis. Unpaired, two-sided Student’s *t*-test was used for comparison of 2 groups. Mantel-Cox log-rank test was used for statistical significance of murine surivival curves. Univariate Cox regression analysis was performed to calculate hazard ratio and log-rank P values per KM-Plotter (58) (http://kmplot.com/analysis/) for the human LAD Kaplan-Meier curves.

## Supporting information

Nuclear Fraction Calculator macro

ROI Draw macro

## Acknowledgement

We thank Dr. John D. Minna for valuable discussions, Drs. Rolf Brekken and David Wang for their helpful comments on the manuscript, Nicolas Loof, Kim Nguyen, and Terry Shih for assistance with FACS, and Denise Ramirez of the Whole Brain Microscopy Facility. We also thank John Shelton at UTSW HistoPathology Core and Dr. Cheryl Lewis at UTSW Tissue Management Core for assistance with tissue processing and embedding. This work was supported in part by the National Cancer Institute (P50CA70907, R01CA196851: J.K.; R21 CA208746: J.-W. K.), National Heart, Lung, and Blood Institute (5T32HL098040 to S.K.), American Cancer Society (RSG-16-090-01-TBG: J.K.), American Lung Association (LCD-400239: J.-W.K.), Sidney Kimmel Foundation for Cancer Research (SKF-14-057: J.K.), Lung Cancer Research Foundation (J.K.) and Bonnie J. Addario Lung Cancer Foundation (J.K.).

