## Supplemental Materials for "Stromal Hedgehog pathway activation by IHH suppresses lung adenocarcinoma growth and metastasis by limiting reactive oxygen species"

**Inventory of Supplemental Information**

❖ **Supplemental Data, Methods, and References**

- Supplement Fig. 1
- Supplement Fig. 2
- Supplement Fig. 3
- Supplement Fig. 4
- Supplement Fig. 5
- Supplement Fig. 6
- Supplement Fig. 7
- Supplement Table 1- qPCR primers
- Supplement Table 2 -Murine sgRNA sequences
- Supplement Material and Methods
- Supplement References

❖ **Custom Macros for ImageJ/Fiji**

- *ROI_Draw*
- *Nuclear_Fraction_Calculator*

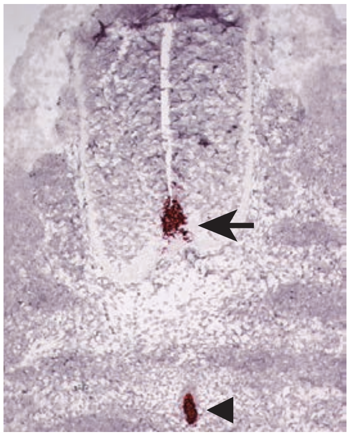

**Supplement Fig. 1** Validation of *Shh* probes for *in situ* hybridization. E11.5 mouse embryonic neural tube is shown. RNA *in situ* hybridization assay was performed with RNAScope using Shh probes. Red puncta indicate *Shh* mRNA. Arrowhead indicates notochord and arrow indicates floor plate.

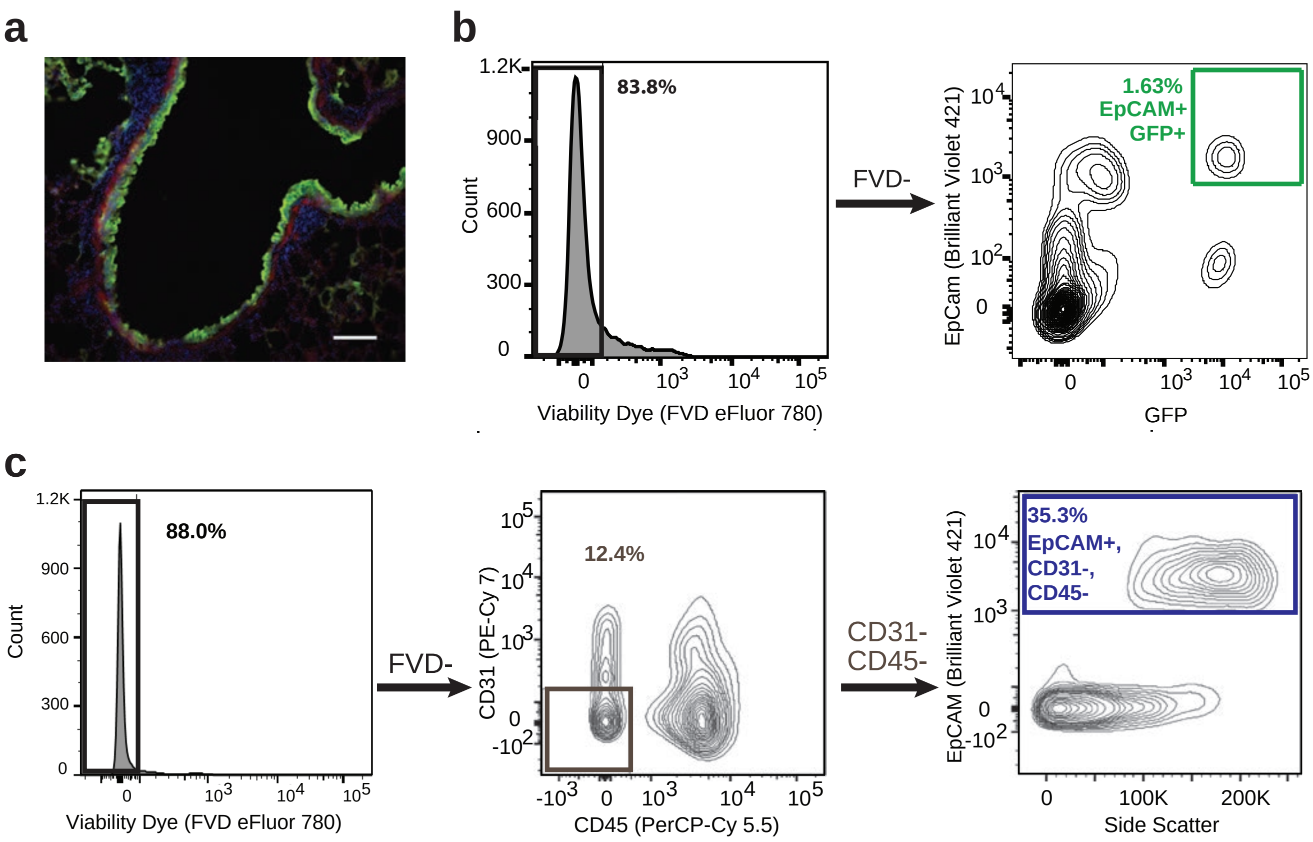


**Supplement Fig. 2** FACS sorting of murine lung epithelial cells. **a** Image of lung section from *KPmTmG* mouse 4 weeks after adeno-cre infection. Adeno-cre infected cells express GFP (green) and uninfected cells express tdTomato (red). Scale bar is 100 micrometers. **b** Lungs of *KPmTmG* mice were subjected to single cell digestion 3 weeks after infection with adeno-cre. Infected lung epithelial cells (EpCAM+,GFP+) were sorted after gating on viable cells (FVD-). **c** Lungs of uninfected *KP* mice were enzymaticcaly digested to single cells and then lung epithelial cells (EpCAM+, CD31-, CD45-) were isloated by FACS after gating on viable (FVD-) and CD31-, CD45- cells.

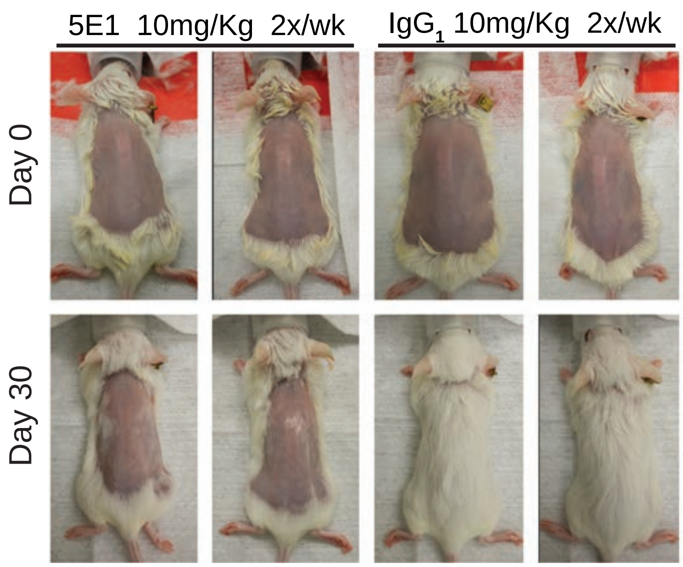


**Supplement Fig. 3** Dose optimization of 5E1 antibody for *in vivo* treatment. Backs of FVB mice were shaved and hair follicles were depilated with Nair to stimulate hair regrowth. 5E1 or $IgG_1$ 10 mg/kg i.p. twice per week were initiated immediately after hair removal for 30 days.

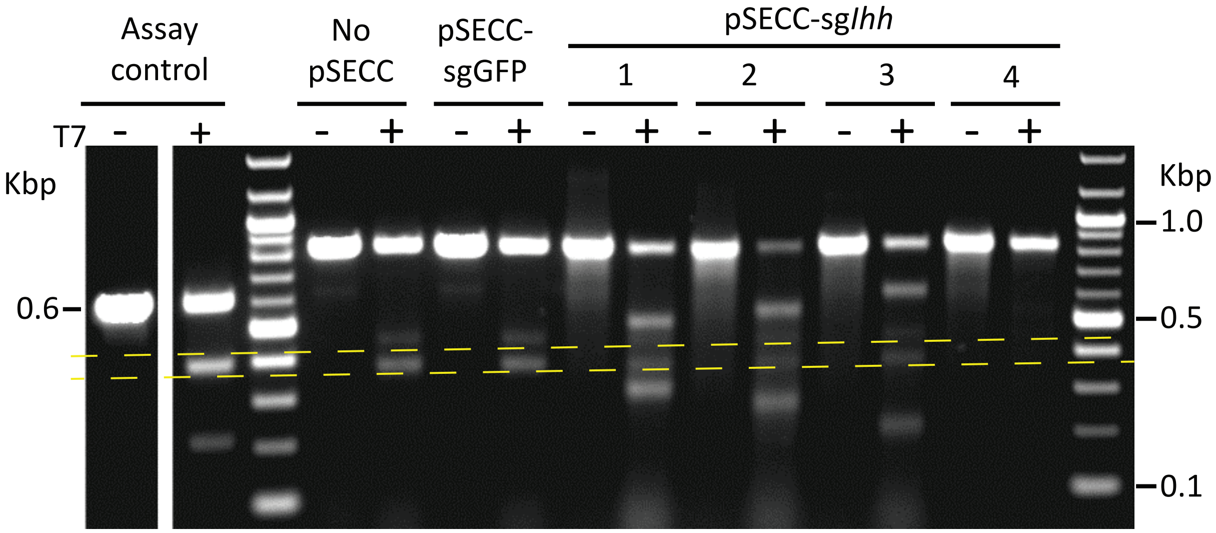


| sgRNA | Expected size of digested products | Percentage of digestion |
|---|---|---|
| GFP | No product | 0.0 |
| 1 | 510 & 320 bp | 44.6 |
| 2 | 550 & 280 bp | 67.4 |
| 3 | 610 & 220 bp | 42.0 |
| 4 | 520 & 310 bp | 0.0 |

**Supplement Fig. 4** SURVEYOR assay of candidate sgRNAs against *Ihh* (sg*Ihh*). Green-Go cells (a reporter cell line that expresses GFP after exposure to cre-recombinase) were infected by lentiviral pSECC-Ihh or pSECC-GFP (negative control). Infected GFP+ cells were isolated by FACS, the genomic target region was amplified by RT-PCR, and subjected to SURVEYOR assay. Bands between the two yellow dashed lines represent non-specific digested products. The table lists the expected size of the digestion products and the percent of digested bands relative to the total RT- PCR product (digested + undigested bands) for each sgRNA based on gel densitometry.

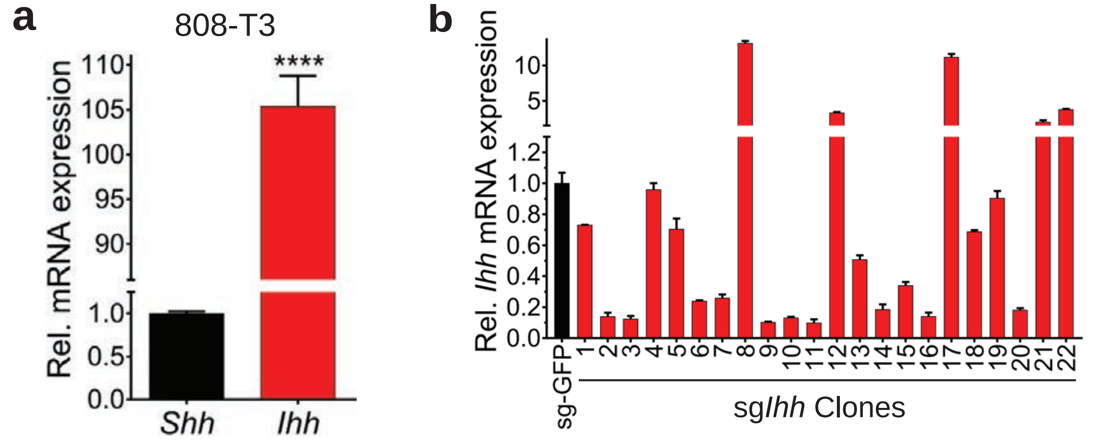


**Supplement Fig. 5** Expression of *Ihh* in KP lung adenocarcinoma cell line. **a** *Shh* and *Ihh* mRNA levels from 808-T3 KP lung adenocarcinoma cells are shown as measured by qPCR. Data are mean of triplicates +/- s.e.m. ****P< 0.0001. **b** *Ihh* mRNA levels of clonal colonies of 808-T3 cells are shown after transient co-transfection with pSECC plasmid containing sgRNA against *Ihh* (pSECC-sg*Ihh*) or GFP (pSECC-sgGFP) and pCMV:DsRed(FRT)GFP plasmid expressing DsRed. DsRed+ transfected cells were FACS sorted and grown as colonies after being plated at limiting dilutions. *Ihh* mRNA levels were measured by qPCR. Bulk pSECC-sgGFP transfected cells were used as control.

| | KRAS mutation | TP53 mutation | IHH ACDbio score | % Acinar | % Lepidic | % Papillary | % Micropapillary | % Solid | Mucinous |
|---|---|---|---|---|---|---|---|---|---|
| Case 1 | G13C | R248L | 0 | 45 | 5 | 0 | 0 | 50 | No |
| Case 2 | **G12A** | **C275F** | **2+** | **10** | **70** | **0** | **5** | **15** | **Yes** |
| Case 3 | **G12V** | **Y220C** | **1+, focal 2+** | **30** | **70** | **0** | **0** | **0** | **Yes** |
| Case 4 | WT | WT | 0 | 30 | 65 | 0 | 5 | 0 | No |
| Case 5 | WT | WT | 0 | 60 | 30 | 5 | 5 | 0 | No |
| Case 6 | WT | WT | 0 | 60 | 30 | 10 | 0 | 0 | No |
| Case 7 | **WT** | **WT** | **1+** | **30** | **70** | **0** | **0** | **0** | **Yes** |
| Case 8 | WT | WT | 0 | 30 | 70 | 0 | 0 | 0 | No |
| Case 9 | WT | WT | 0 | 5 | 5 | 30 | 10 | 50 | No |

**Supplement Fig. 6** Summary of IHH mRNA *in situ* hybridization and histological features of human lung adenocarcinoma samples. Data corresponds to Figure 5A. Samples that are positive for IHH mRNA are in bold.

| Median Survival (Days) | | | |
|---|---|---|---|
| 5E1 | | $IgG_1$ | |
| Control | 88 | Control | 99 |
| NAC | 97 | NAC | 96 |

**Supplement Fig. 7** Median survival of mice from Figure 6f and g. Median survival of *KP* mice treated with 5E1 and NAC (Fig. 6f) or IgG1 with NAC (Fig. 6g).

| Gene | Forward Primer (5' → 3') | Reverse Primer (5' → 3') |
|---|---|---|
| Human *HPRT1* | GGTCAGGCAGTATAATCCAAAG | GGACTCCAGATGTTTCCAAAC |
| Mouse *Gapdh* | CATTTGCAGTGGCAAAGTGGAG | ACCCCATTTGATGTTAGTGGGG |
| Human *SHH* | CAAGCAGTTTATCCCCAATGTG | TCACCCGCAGTTTCACTC |
| Mouse *Shh* | CTATGAGGGTCGAGCAGTGG | GAAACAGCCGCCGGATTTG |
| Human *IHH* | CCGGCTTTGACTGGGTGTAT | ATGAGCACATCGCTGAAGGT |
| Mouse *Ihh* | CCAGTGGCCTGGTGTGAAAC | CGTGGGCCTTGGACTCGTAA |

Supplement Table 1. qPCR primers used in this study.

| sgRNA | Sequences (5' → 3') |
|---|---|
| *GFP* [ref. (1)] | GGCCACAAGTTCAGCGTGTC |
| *Ihh*-1 | CTGGGTGTATTACGAGTCCA |
| *Ihh*-2 | ACTGCTGGCGCGCTTAGCAG |
| *Ihh*-3 | TTTACACTATGAGGGCCGCG |
| *Ihh*-4 | GTAATACACCCAGTCGAAGC |

Supplement Table 2. Murine sgRNA sequences used in this study.

### Supplemental Material and Methods

#### 5E1 generation and purification

5E1 Hybridoma cells (Developmental Studies Hybridoma Bank) were maintained in Hybridoma medium (Gibco) with 20% FBS (heat inactivated super low IgG serum, HyClone), 1X GlutaMax (Gibco), and 2.5 mM HEPES (Fisher Scientific). Cells were maintained for 7 days and then supernatant was collected, centrifuged, and filtered through a 0.2 μm filter. Presence of 5E1 in the supernatant was verified with an ELISA assay. 3M NaCl and 0.1M Borate were added to the supernatant and pH adjusted to 8.5 and loaded on to a protein A bead column using a peristaltic pump at 1.5 mL/min at 4°C for 3 rounds. 5E1 was eluted using ImmunoPure Genetle Ag/Ab Elution Buffer (PIERCE) and then subjected to dialysis in PBS using Slide-A-Lyzer Dialysis Cassette (Thermo Fischer Scientific).

#### RNA in situ hybridization analysis of human lung adenocarcinoma samples

RNA *In situ* hybridization (ISH) was conducted by automated RNAscope assay using the Leica Bond RX autostainer (Leica Biosystems, Nussloch, GmbH) to visualize single RNA molecule per cell as a single dot in FFPE tissue samples at least. The procedure performed on BOND RX system are briefly described as the following. Tissue sections of 5um thick were deparaffinized and rehydrated, then followed with the Leica Bond prestaining and staining protocols. Antigen retrieval was performed with ER2 (BOND Epitope Retrieval Solution 2) at 95°C for 15 minutes and ACD Protease treatment at 40°C for 15 min; then RNA-specific probes were hybridized to target RNA for 120 min. The signals were amplified by multiple steps, which was followed by hybridization to horseradish peroxidase (HRP)-labeled probes and detection using the 3,3′-Diaminobenzidine (DAB) chromogenic substrate. After that, cell nuclei were counterstained with hematoxylin. The RNAscope probes assayed in this study were: IHH (target probe) (RNAscope® LS 2.5 Probe- Hs-IHH, 472388, Advanced Cell

Diagnostic, Hayward, CA, USA), PPIB (housekeeping gene, positive control probe) (Hs-PPIB, 313908, Advanced Cell Diagnostic, Hayward, CA, USA) and dapB (bacterial gene, negative control probe) (312038, Advanced Cell Diagnostic, Hayward, CA, USA). Positive and negative control probes were used to assess RNA quality of tissue section and optimal permeabilization.

RNA expression of IHH was scored using conventional bright-field microscopy in adenocarcinoma malignant cells. We applied a semi-quantitatively scoring system, in a scale of 0-4, based on the number of RNA copies per cell (0, no staining or <1 dot/10 cells; 1+, 1-3 dots/cell; 2+, 4-9 dots per cell, None or very few dot clusters; 3+, 10-15 dots/ cell and <10% dots are in clusters; 4+, >15 dots/cell and >10% dots are in clusters,). Positive (PPIB) and negative (Dapb) control probes were also evaluated, dapB score of <1 and PPIB score ≥2 with relatively uniform PPIB signals throughout the sample were considered adequate for analysis (data not shown).

**Lentivirus generation**

Lentivirus containing pSECC-Ihh and pSECC-GFP were generated as described previously with some modifications (2). Briefly, 85-90% confluent 293-T cells in 15 cm cell culture dish were co-transfected with 16 μg pSECC-*Ihh* or pSECC-GFP, 8 μg pCMV-dR8.91 (packaging vector), and 4 μg pCMV-VSV-G (enveloping vector) using Lipofectamine 3000 (Thermo Fischer Scientific). pCMV-dR8.91 and pCMV-VSV-G vectors were kindly provided by Dr. John Minna (UTSW). Growth medium was replaced after 16 hours, subsequent supernatant was collected 48 hours later and centrifuged at 800xg at 4°C to precipitate cell debris. Supernatant above the cell debris pellet was collected and ultracentrifuged at 100,000xg for 2 hours.

Subsequent pellet containing the lentivirus was resuspended in OptiMEM (Gibco). Titration of lentivirus was performed per previously described protocol (https://tinyurl.com/yd2qk9l6).

*SURVEYOR assay***.** Lentivirus containing selected pSECC-*Ihh* was generated in a small scale (10 cm cell culture plate) using 293T cells as described above. Cell culture supernatant (containing lentivirus) was collected 72 hours after transfection and placed on 10 cm cell culture dish of 70-80% confluent Green-Go reporter cells and maintained for 48 hours. GFP+ Green-Go cells (2) were isolated by FACS. Genomic DNA from GFP+ cells was isolated using PureLink Genomic DNA Mini Kit (ThermoFischer Scientific). The sg-*Ihh* target region of genomic DNA was amplified by PCR using the following primers: forward 5'-GTCCCATGAGTGCTGTCGA-3' and reverse 5'-TGACCTGCATTTGCGTGGTA-3'. Mutation detection on PCR product was performed using EnGen Mutation Detection Kit (New England BioLabs) and following manufacturer instructions.

**Analysis of γH2AX+ nuclei in immunohistochemical stained lung sections**

Slides with immunohistochemically stained lung sections were scanned at 40x magnification using a Hamamatsu Nanozoomer. The scanned images were exported as tiff images at 10x resolution. Analysis was performed on Fiji imaging software. Tumors were outlined interactively using the custom "ROI_Draw" macro. To obtain the fraction of γH2AX positive nuclei in tumors, a custom macro, "Nuclear Fraction Calculator", was used. Briefly, the "Nuclear_Fraction_Calculator" macro operates as follows. DAB positive nuclei are segmented using the "Colour Deconvolution" plugin written by Gabriel Landini (Image>Color>Colour Deconvolution in Fiji or as an ImageJ plugin available from (https://tinyurl.com/yckvnp3f). The built-in H DAB matrix is used to separate DAB staining from hematoxylin staining, the DAB channel is converted to 8-bit grayscale, the contrast is inverted, and a binary mask is generated by thresholding the image using the maximum entropy method, giving the user an option to adjust the threshold interactively. A watershed operation is used to separate connected nuclei and then ultimate points is used to represent each object with a dot. To segment the total nuclei, the original color image is converted to 8-bit grayscale and the “Mexican Hat Filter” plugin by Dimiter Prodanov (https://tinyurl.com/y7vmzrow ) was used to smooth the image and enhance the edges of the nuclei. A binary mask is made, followed by watershed and ultimate points to mark each object. The previously stored ROI set is used together with these masks for analysis of each tumor to obtain the area of the tumor, the number of DAB stained (γH2AX+) nuclei, the total number of nuclei and the fraction of DAB stained (γH2AX+) nuclei. “ROI_Draw” and “Nuclear_Fraction_Calculator” macros are compatible with ImageJ and Fiji.
