## Supplementary material for "Stromal Hedgehog pathway activation by IHH suppresses lung adenocarcinoma growth and metastasis by limiting reactive oxygen species": Nuclear Fraction Calculator macro

//This macro was developed using ImageJ 1.52e with 64-bit Java 6 JRE. For any questions or concerns contact Dr. Kate Phelps at. 
//Open the image and then run the macro.
//The macro finds DAB positive nuclei in slide scanner images of immunohistochemically stained tissue sections counter-stained with hematoxylin.
//Additional plugins required for this macro:
	//Colour Deconvolution by Gabriel Landini (included in Fiji under the Image>Color menu) or available from GitHub (https://github.com/fiji/Colour_Deconvolution/releases/tag/Colour_Deconvolution-3.0.2)  
	//Mexican Hat Filter plugin developed by Dimiter Prodanov. Available here: https://imagej.nih.gov/ij/plugins/mexican-hat/index.html

//First it checks to see if an image is open - the macro exits if no image is open

if (nImages==0) 
exit("No images are open. Please open an image and restart the macro");

//The macro assumes that regions of interest have already been drawn and saved as a collection named "imagefilename _ROISet.zip"

//Next, the macro clears out the roiManager and declare arrays to store nuclei counts and area for each region of interest. The array length will equal the number of regions in the ..ROISet.zip file for this image. The results for each region will be output in the results table at the end.

roiManager("reset");
AreaArray=newArray();
KiNArray=newArray();
TotalArray=newArray();
PercentArray=newArray();

//Get name of image and directory path. This information will be used for labeling results and saving files.

filename=getTitle;
dir=getDirectory("image");

//Optionally hide image windows. Comment the next line out to stop hiding windows.

setBatchMode(true);

//Generate three working images called Original, DAB+ Mask and Total Nuclei Mask 

showStatus("Creating working images...");

//rename the original image to Original.
rename ("Original");
run("Select None");

//DAB+ Mask is generated by color deconvolution of the H DAB image and then converting the DAB channel to 8-bit grayscale and inverting the contrast.
run("Colour Deconvolution", "vectors=[H DAB]");
selectWindow("Original-(Colour_1)");
close();
selectWindow("Original-(Colour_3)");
close();
selectWindow("Colour Deconvolution");
close();
selectWindow("Original-(Colour_2)");
run("8-bit");
run("Invert");
rename("DAB+ Mask");

//Total Nuclei Mask is generated from Original and converted to 8-bit grayscale
selectWindow("Original");
run("Duplicate...", "title=[Total Nuclei Mask]");
run("8-bit");

//Segment the two masks to produce binary images showing the location of each nucleus; the user has the option to adjust the threshold used to segment the DAB+ Mask.
//Resulting images are saved in the same folder as the original file

showStatus("Segmenting total nuclei...");
wait(5);

selectWindow("Total Nuclei Mask");
run("Mexican Hat Filter", "radius=5");
run("Make Binary");
run("Watershed");
run("Ultimate Points");
setThreshold(1, 255);
run("Convert to Mask");
save(dir+"/"+filename+"_Total Nuclei Mask");

showStatus("Segmenting DAB+ nuclei...");
wait(5);
selectWindow("DAB+ Mask");
run("Threshold...");
setAutoThreshold("MaxEntropy dark");
selectWindow("Threshold");
setBatchMode("show");
waitForUser("Please adjust the threshold so only the DAB+ nuclei are red (Zoom in and use the original image as a guide");
setBatchMode("hide");
run("Convert to Mask");
run("Watershed");
run("Ultimate Points");
setThreshold(1, 255);
run("Convert to Mask");
save(dir+"/"+filename+"_DAB+ Mask");


//Open the saved ROISet file associated with this image and get area of each ROI

roiManager("Open",dir+"/"+filename+"_ROISet.zip");
count=roiManager("count");
showStatus("Counting nuclei...");
for (k=0;k<count;k++){
index=k+1;
selectWindow("Original");
run("Select None");
roiManager("Select", k);
roiManager("measure");
roiArea=getResult("Area");
AreaArray=Array.concat(AreaArray,roiArea);
}

//Count DAB+ nuclei and save as ROI set (color of these set to red). 
//A separate set of DAB+ nuclei will be saved for each ROI analyzed. These can be opened later and used to check how well the macro performed.

for (k=0;k<count;k++){
index=k+1;
selectWindow("DAB+ Mask");
run("Select None");
roiManager("Select", k)
run("Analyze Particles...", "  show=Nothing display exclude clear add");
roiManager("Combine");
roiManager("Add");
lastK=roiManager("count");
lastK=lastK-1;
roiManager("Set Color", "red");
roiManager("Set Line Width", 0);
roiManager("Select", lastK);
roiManager("save", dir+"/"+filename+"_DAB+ nuclei_Region"+index+".zip");
KiN=nResults;

//Count total nuclei and save as ROI set. Color of these will be  yellow. A separate set of Total Nuclei will be saved for each ROI analyzed.

selectWindow("Total Nuclei Mask");
roiManager("reset");
run("Select None");
roiManager("Open",dir+"/"+filename+"_ROISet.zip");
roiManager("Select", k);
run("Analyze Particles...", "  show=Nothing display exclude clear add");
roiManager("Combine");
roiManager("Add");
last=roiManager("count");
last=last-1;
roiManager("Set Color", "yellow");
roiManager("select", last);
roiManager("save", dir+"/"+filename+"_Total nuclei_Region"+index+".zip");
total=nResults;

//Calculate the % DAB+ nuclei

KiNpercent=KiN*100/total;

//Store the number of DAB+ nuclei, total nuclei and % DAB nuclei each in their own array


KiNArray=Array.concat(KiNArray,KiN);
TotalArray=Array.concat(TotalArray,total);
PercentArray=Array.concat(PercentArray,KiNpercent);

roiManager("reset");
roiManager("Open",dir+"/"+filename+"_ROISet.zip");
}

//Output the results in a table

run("Clear Results");
length=KiNArray.length;
for (n=0;n<length;n++){
setResult("Image Name",n,filename);
setResult("Region #",n,n+1);
setResult("Area in pixels",n,AreaArray[n]);
setResult("DAB pos",n,KiNArray[n]);
setResult("Total nuclei",n,TotalArray[n]);
setResult("% DAB pos",n,PercentArray[n]);
}

showStatus("Anaysis Complete");
