## Supplementary material for "Stromal Hedgehog pathway activation by IHH suppresses lung adenocarcinoma growth and metastasis by limiting reactive oxygen species": ROI Draw macro

//Get name of image and directory path. For labeling results and saving files. For any questions or concerns contact Dr. Kate Phelps at. 

filename=getTitle;
dir=getDirectory("image");
roiManager("reset");
if (File.exists(dir+"/"+filename+"_ROISet.zip"))
roiManager("Open",dir+"/"+filename+"_ROISet.zip");
roiManager("show all with labels");

//Draw region(s) of interest outlining each tumor; to combine multiple outlines hold down shift. The ROIs will be combined and saved in one file.

setTool("freehand");
Ans=1;
while (Ans==1){
waitForUser("Please use the Freehand Tool to outline the tumor; Click OK when done");
roiManager("Add");
count=roiManager("count");
roiManager("Select",count-1);
roiManager("rename","Region"+count);
roiManager("save", dir+"/"+filename+"_ROISet.zip");
Ans=getBoolean("Draw another ROI?");//Clicking no will stop the loop and move to the next line of the macro
}
